# Neutral mutation disequilibrium is common in prokaryotes and animals

**DOI:** 10.1101/2025.08.22.671660

**Authors:** Katherine Caley, Benjamin D. Kaehler, Von Bing Yap, Gavin A. Huttley

## Abstract

Widely used models of sequence evolution assume DNA sequences are at equilibrium with respect to the processes that drive divergence, mutation and natural selection. Violations of this assumption have been shown to affect the conclusions from analyses of genetic variation. However, there is currently a lack of statistical methods to assess whether this phenomenon occurs with sufficient frequency and magnitude to warrant concern. Here, we introduce such techniques and apply them to cases where genetically encoded changes to mutation are expected to induce neutral mutation disequilibrium (NMD) across an entire genome (loss of the hypermutable base 5-methyl-cytosine in *Drosophila melanogaster*), or in a limited genomic region (translocation of the *Fxy* gene to the pseudo-autosomal region in *Mus musculus*). We find that the proportion of 9,237 genes in NMD in the *D. melanogaster* genome (∼0.81) is markedly elevated compared to its sister taxa *D. simulans* (∼0.31) which retains 5-methyl-cytosine. We further establish that the translocation of *Fxy* has resulted in a ∼ 3× increase in the magnitude of NMD. These empirical case studies validate the robustness of our methods. Applying the methods to human evolution, from 613 genes, we estimated the proportion of NMD for the exons as ∼0.64 and introns as ∼0.22. These results establish NMD as a common phenomenon and suggest that conclusions from studies that assumed equilibrium in order to understand biological mechanisms may misattribute to biology what should be credited to measurement error.

**Significance:** It has been famously said that all models are wrong, but some are useful. Statistical models of sequence divergence provide the fundamental quantities from which we derive meaning about the causative mechanisms responsible for patterns in molecular evolution. Published work has established current models are not just wrong, but misleading – incorrectly assuming neutral mutation equilibrium leads to errors in the inference of the mode of natural selection and the existence of the molecular clock. In this work, we demonstrate that neutral mutation disequilibrium is common, suggesting that these errors may be widespread. Our results raise the strong likelihood that detected patterns in molecular evolution are being misattributed to biological mechanisms when, in fact, they reflect systematic measurement error.

## Introduction

The vast majority of statistical methods concerned with biological sequence data embed a mathematical representation of mutation. Such methods underpin a wide range of applications, including being used to characterize the mutation spectra of cancers (e.g. Nik-Zainal et al. 2012), estimate the neighborhood effects on point mutation (e.g. Zhu et al. 2017), identify regions subjected to evolutionary constraint, or model the substitution process for reconstructing phylogenies (Felsenstein 1981). Most pervasively in the phylogenetic reconstruction case, substitutions are assumed to originate from processes that are stationary (frequencies of nucleotides do not change through time) and time-reversible (identical whether run in forward or reverse). Stationarity requires a state of mutation equilibrium which is almost impossible if the processes affecting mutagenesis change through time. This makes it a particularly troubling assumption given the overwhelming evidence of heritable differences in DNA metabolism (Sampson et al. 2015), and thus mutagenesis. With advances in whole genome DNA sequencing, empirical mutagenesis studies have indeed indicated that neutral mutagenesis does not remain stable through time (Long et al. 2018; Sane et al. 2023). Here, we introduce statistical techniques that complement the empirical data by exploiting signal from the operation of mutagenic processes over millions of years. In doing so, we answer the question of how pervasive neutral mutation disequilibrium is.

We consider a DNA sequence to be at “mutation equilibrium” if its nucleotide frequencies remain the same through time. The case of disequilibrium is thus when this condition does not hold. If the divergence process is strictly in accordance with mutation equilibrium then, from the Neutral Theory (Kimura 1983), the composition of related sequences should remain largely unchanged. Observations of substantial differences in nucleotide frequency between genomes (e.g. Karlin 1998) imply the existence of historical mutation disequilibrium. We will refer to these as neutral mutation equilibrium (NME) and neutral mutation disequilibrium (NMD) respectively. Empirical mutagenesis studies have confirmed a striking capacity for variability in mutagenic processes (e.g. Kondrashov 2008). Studies of *de novo* mutation in parent-offspring trios provide strong support for substantial genomic heterogeneity in mutation processes (Michaelson et al. 2012; Francioli et al. 2015; Smith et al. 2018; Bergeron et al. 2023). Mutation accumulation (MA) experiments in cell lines have shown that lineage-specific mutational biases shape nucleotide composition (Long et al. 2018) and even suggest that shifts reversing the prevailing bias may be favored by selection (Sane et al. 2023; Tuffaha et al. 2023). Together, these findings indicate that real divergence processes routinely change, putatively violating the widespread assumption of NME.

An alternate approach to interrogating mutational processes is to exploit the accumulation of genetic changes between species using non-stationary models of sequence evolution. Squartini and Arndt (2008) developed a hypothesis test and measures of the magnitude of non-stationarity based on a non-stationary strand-symmetric substitution process. Application of their methods to divergence within Drosophila (Singh et al. 2009; Squartini and Arndt 2008) was used to establish substantial inter-lineage heterogeneity in substitution processes, the existence of NMD and its differences between autosomes and the X-chromosome.

A desirable benchmark for statistical methods is data from a natural system where the signal of interest should exist. Changes to mutagenesis are expected to elevate NMD while the nucleotide frequencies equilibrate to the new mutagenic process. Therefore, by contrasting a sample with perturbed mutagenesis with a corresponding unperturbed sample, one can predict directional inequalities that statistical methods intended to capture NMD should reproduce. At the time of the works described above, there were no widely accepted empirical cases that met this criteria, precluding such empirical validation. We now summarize two cases that do satisfy these characteristics that we employ in this study as empirical positive controls.

The recent loss of DNA methylation in *Drosophila melanogaster* is expected to have elevated NMD across its genome. Since divergence from its common ancestor with *D. simulans, D. melanogaster* has lost multiple components of the enzymatic machinery concerning 5-methyl-cytosine (^5^mC) (Lyko 2018) and exhibits markedly reduced levels of ^5^mC compared to *D. simulans* (Capuano et al. 2014; Deshmukh et al. 2018). This relative depletion of the hyper-mutagenic ^5^mC (Coulondre et al. 1978) will have induced a major shift of the mutagenesis spectrum in *D. melanogaster*, leading to the prediction of a genome wide elevation of NMD relative to *D. simulans*.

The evolutionary history of the *Fxy* gene in *Mus musculus* indicates that it too has been subjected to a substantial change in mutagenesis. In *M. musculus, Fxy* has been translocated from a strictly X-linked location to a location where it straddles the boundary of the Pseudo-Autosomal Region (PAR, Figure S1) (Palmer et al. 1997). The PAR is characterized by elevated rates of meiotic recombination between the X and Y chromosomes. Meiotic recombination, and its association with GC-biased gene conversion, has been implicated as a driver of regional base composition differences (Montoya-Burgos et al. 2003). Additionally, PAR-linked sequences are subjected to distinctive male germline mutation processes (e.g. Huttley, Jakobsen, et al. 2000). Based on these properties, we hypothesized that there should be a localized elevation of NMD in the portion of *Fxy* located in the PAR compared to that which remains X-linked.

The observation that real homologous nucleotide sequences frequently exhibit compositional differences has motivated the development of models which relax the assumptions of time-reversibility and stationarity. It is important to note that time-reversibility implies stationarity, but a stationary process need not be time-reversible. Stationary non-time-reversible models (Yang 1994; Yang and Roberts 1995), and the more general non-stationary non-time-reversible models (Yap and Speed 2005) were introduced some time ago. However, software implementation of these models suitable for exploratory analysis is relatively recent (Kaehler, Yap, Zhang, et al. 2015). The work of Kaehler, Yap, Zhang, et al. 2015 also provided the first systematic assessment comparing a non-stationary process to competing time-reversible models. Their results demonstrated that non-stationary models fit the empirical data substantially better than time-reversible models. Further, their demonstration of systematic errors in genetic distance estimates under the assumption of stationarity suggested that non-stationary processes are prevalent in natural data. Here we directly evaluate that possibility.

We have sought to develop the statistical procedures to determine how widespread NMD is and to measure its magnitude. In doing so, we benchmarked performance of our methods against the expectations for the empirical datasets articulated above: an elevated occurrence and magnitude of NMD in *D. melanogaster* compared to *D. simulans*; and, a higher NMD magnitude in the segments of *Fxy* recently translocated into the PAR compared to their X-linked counterparts. We show that the statistics capture these expected phenomena very well, successfully identifying global and local NMD. These results support the credibility of the estimated NMD frequency within the human genome, lending support to the argument that NMD may be taxonomically widespread. Within both the empirical knowns and the human case, more granular features of the decay of NMD reflecting the influence of natural selection on this phenomenon are discussed.

## Results

We have developed a formal hypothesis test for the occurrence of NMD (Test of Existence, TOE) and a measure of NMD magnitude (∇) by analysis of synthetic data with properties we expect to be challenging (TOE and ∇ defined in Methods, see Figure 1 for an overview). Nucleotide frequencies are integral to the definition of NMD and we furthermore want to avoid confounding from historic NMD, whereby sequences differ in their composition even though they have reached equilibrium. We therefore considered proxy measures of both of these properties and devised experiments to establish that our statistical procedures are robust to combinations of their extremes. The presence or absence of historic NMD between taxa was quantified using Jensen-Shannon divergence (JSD). Whether the nucleotide frequency distribution was balanced or imbalanced was quantified using Shannon’s entropy. From published analyses of taxonomic triples of microbial 16S rRNA sequences, we identified four alignments (Figure S3) that represented the different combinations of Low or High JSD and Low or High Shannon’s entropy. Using the null model of the TOE (Figure 1a) as the generating process, these alignments were used as seeds for synthetic data sets. We further considered the impact of sampling error using systematic variation in the alignment length.

**Figure 1.**
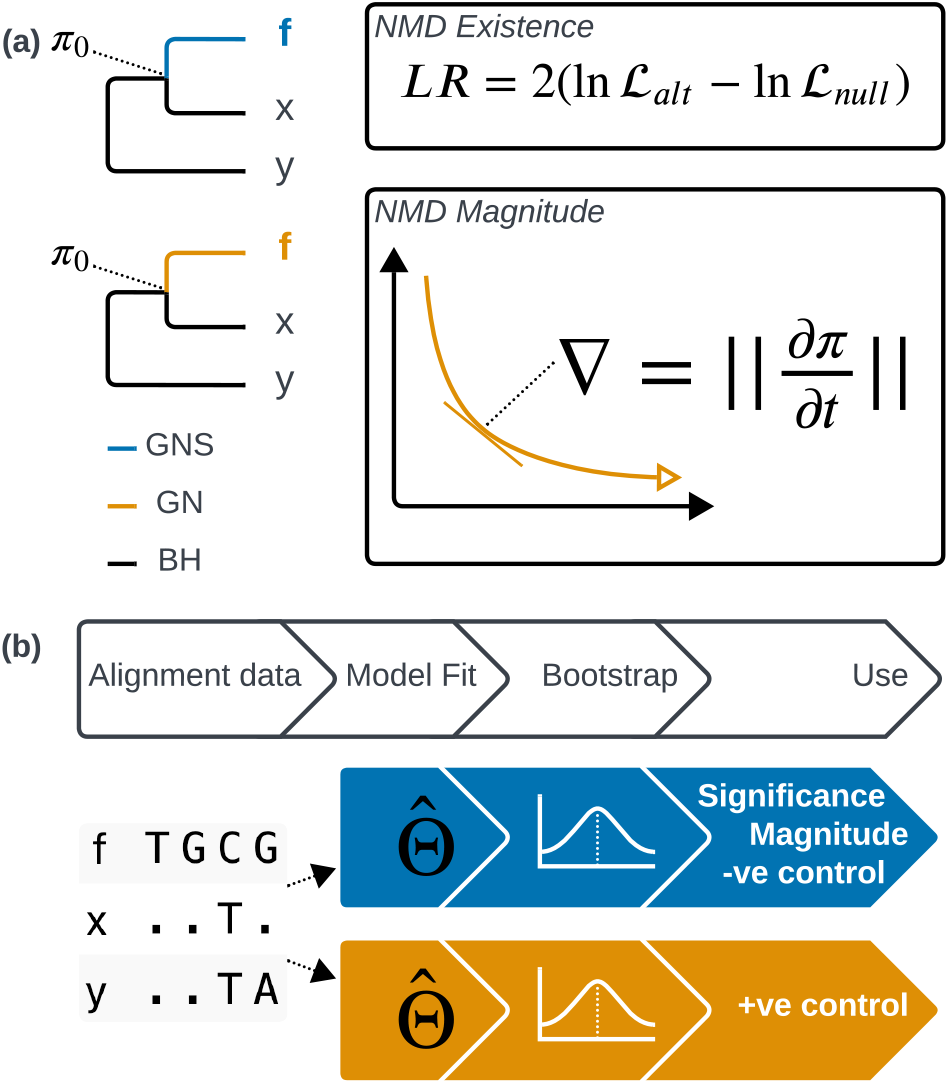
Statistics and analyses for facilitating robust inference of neutral mutation disequilibrium (NMD). (a) Statistics used to quantify NMD. The branch to **f** is designated the foreground branch, and the remainder are background branches. The Markov process applied to substitutions on **f** distinguishes the null and alternate hypotheses: the non-reversible, stationary (GNS) Markov process is used for the null with *π*_0_ = *π*_∞_ for **f**; the non-reversible, non-stationary (GN) Markov process is used for the alternate with no constraints on *π*_0_. Uses of these models are identified by colour. Substitutions on the background branches are always represented using the Barry Hartigan model (BH, Barry and Hartigan 1987) Markov process. The LR of these two hypotheses is used to test for the existence of NMD. The magnitude of NMD on **f** is the rate of change in nucleotide frequency using maximum likelihood estimates (MLEs) obtained from the alternate (denoted 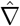 , see Equation 1) when the process is sampled at the dotted line. (b) Parametric bootstraps are applied to adjust NMD statistics and generate control samples to guide inference. MLEs from the null model are used to estimate the *p*-value, correct the magnitude estimate, and produce negative controls. MLEs from the alternate model are used to produce positive controls.

### Simulation Studies Demonstrate That Robust Detection and Quantification of NMD Requires Parametric Bootstraps

We examined the behavior of the TOE and ∇ in data generated under the null. For such data, where the generating process is in NME, we expect the following conditions to hold. (1) *p*-values from the TOE should be uniformly distributed; and (2) our measure of NMD magnitude, 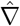, should approach zero.

Applying the TOE to the synthetic datasets established that parametric bootstrapping was required to derive reliable *p*-values. As demonstrated in Figure 2, for all alignment lengths there was a tendency to under-estimate small probabilities (Figure 2a) across all simulation conditions (Figure S4) which would lead to a higher false discovery rate. In general, as alignment length increased the distribution of *p*-values approached the theoretical expectation. However, even for the longest alignments (30,000 bp), the distributions were not uniform.

**Figure 2.**
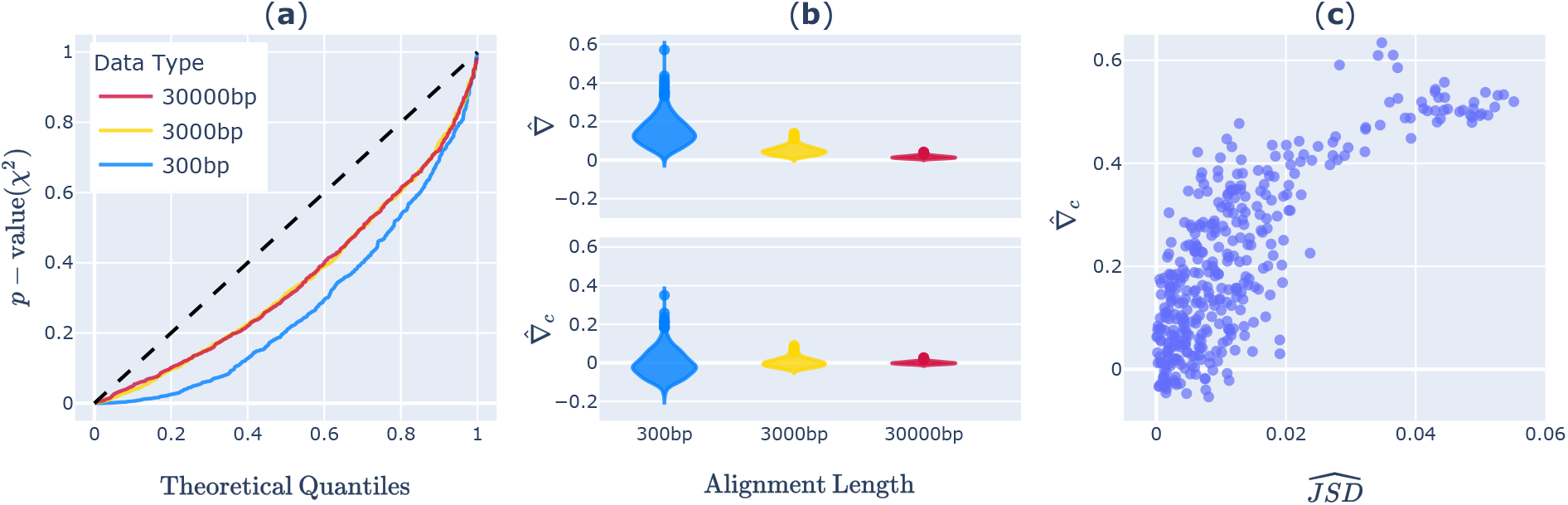
Properties of measures of Neutral Mutation Disequilibrium from the High JSD and High Entropy condition. When the null hypothesis is true (a) *p*-values from the test-of-existence (derived from the *χ*^2^ distribution) for mutation disequilibrium do not match theoretical expectation shown as the dashed diagonal (see Figure S4 for the quantiles for the other conditions), (b) the magnitude of mutation disequilibrium captured by 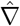 was sensitive to alignment length, which was corrected for by 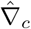. For observed microbial 16S rRNA sequence data (c) 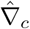 increased with increasing JSD.

Application of parametric bootstrapping was required to correct our estimates of NMD magnitude for the positive bias of 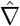 that was inversely proportional to the alignment length. As a Euclidean distance, 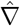 is always positive and thus was expected to reflect both NMD and sampling error. For alignment lengths typical of molecular evolutionary analyses, the sampling errors were pronounced (Figure 2b). We therefore introduced a corrected measure, ∇_*c*_, (Equation 2) which subtracts the mean of 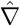 from parametric bootstraps under the null. As shown in Figure 2b, this correction was effective in centering the statistic on 0, removing the bias due to alignment length. Finally, we confirmed in empirical data that the predicted positive correlation exists between our measure of NMD magnitude 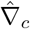 and the NMD proxy measure JSD (Figure 2c).

### The Translocation of *Fxy* into the Pseudo-Autosomal Region (PAR) Validates Localized Detection of NMD

Our analysis of *Fxy* provided support for the hypothesis that local genomic rearrangements induce measurable NMD. PAR-linked introns exhibited a higher level of NMD than their X-linked counterpart (Table 1). Four alignments of *Fxy* introns were found suitable for analysis (see Methods). In *M. musculus*, intron rank 2 remains X-linked while ranks 4-6 reside within the PAR. We rejected the null hypothesis of NME for the PAR-linked introns only, providing evidence for the existence of NMD in precisely the PAR-linked region. Furthermore, estimates of the magnitude of NMD were ∼3× higher for PAR-linked introns compared to that of the X-linked intron (Table 1).

**Table 1:** The magnitude of mutation disequilibrium is higher in the region of the *Fxy* gene within the PAR. Intron rank 2 remains X-linked while ranks 4-6 are within the PAR. 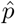-value and 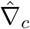 are from a TOE with *M. musculus* treated as the foreground branch, 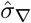 is the estimated standard deviation of from the null distribution, length is from the sampled *M. musculus* intron sequence. A 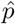-value of 0.00 indicates only that we did not detect a larger LR in the 100 bootstrap replicates.

| Statistic \ Intron rank | 2 | 4 | 5 | 6 |
| --- | --- | --- | --- | --- |
| $\hat{p}$-value | 0.20 | 0.00 | 0.00 | 0.00 |
| $\hat{\nabla}_c$ | 0.24 | 0.73 | 0.74 | 0.66 |
| $\hat{\sigma}_{\nabla}$ | 0.10 | 0.03 | 0.05 | 0.08 |
| length | 3,538 | 4,358 | 1,864 | 834 |

### Genome-wide Elevation of NMD in *D. melanogaster* Following Loss of DNA Methylation

Our analysis of *Drosophila* genomes provided strong support for the hypothesized impact of methylation loss, showing a significant increase in the occurrence of NMD in *D. melanogaster* compared to *D. simulans* (Figure 3). Using the TOE, the estimated proportion of loci supporting the alternate hypothesis (Storey and Tibshirani 2003) of NMD was approximately three-fold greater in *D. melanogaster* (0.83) compared to *D. simulans* (0.31). The distribution of the TOE test statistic is presented using “smile plots” (see Methods), which display the observed distribution alongside those of synthetic positive and negative control datasets (Figure 3). In *D. melanogaster*, the observed data was closest to that of the positive control, consistent with a genome-wide elevation in NMD, while in *D. simulans* the observed data were closer to, but still above, the negative control (Figure 3a-b).

**Figure 3.**
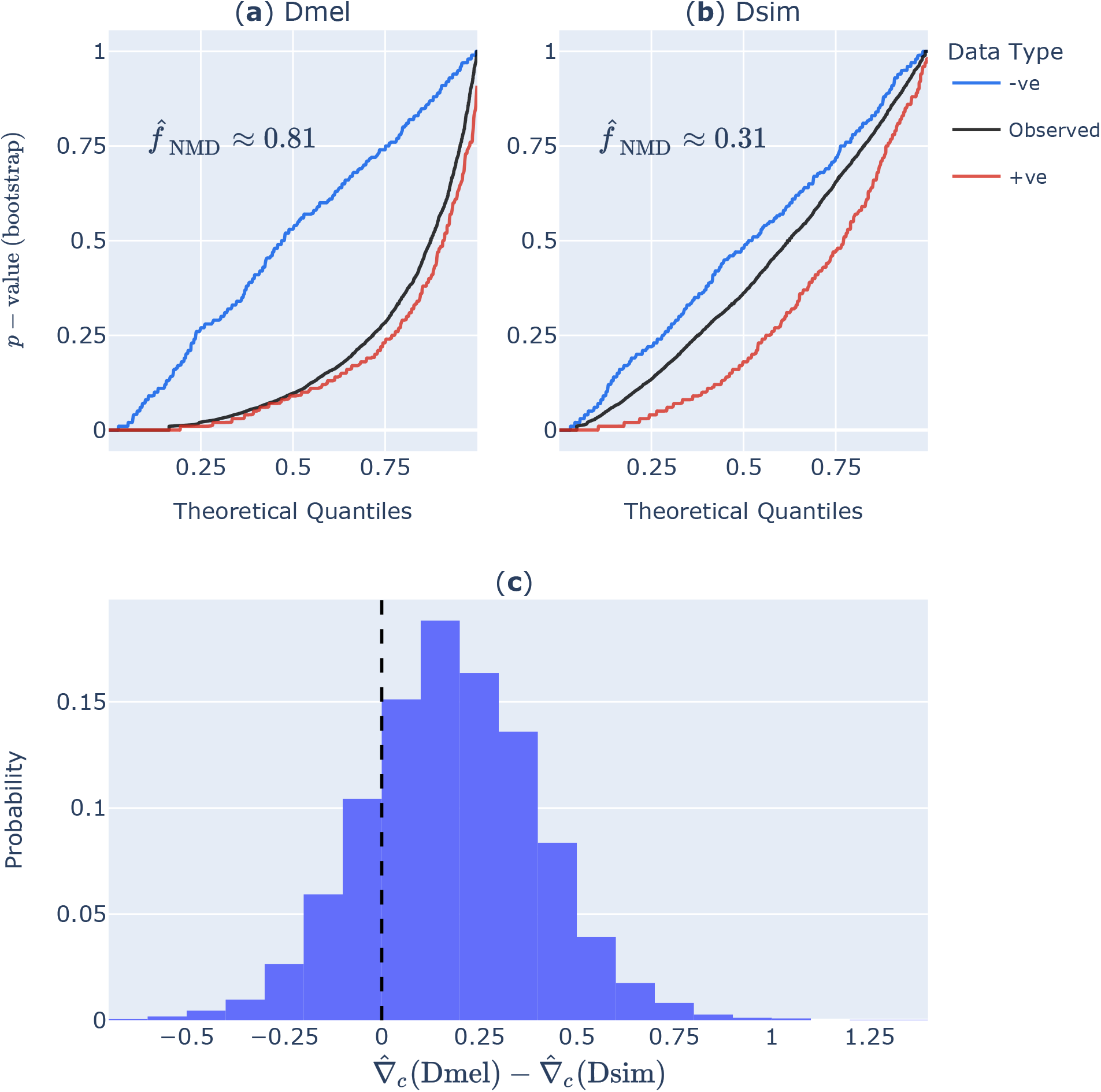
Systematically greater NMD in *D. melanogaster* compared to *D. simulans*. Smile plots showing the quantile distribution of 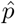-values from the TOE with (a) *D. melanogaster* (Dmel) and (b) *D. simulans* (Dsim) as the foreground branch. 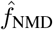-estimate of frequency of alignments in neutral mutation disequilibrium for the respective lineage (Storey and Tibshirani 2003). Data Type indicates the alignment origins with −ve, +ve being the negative and positive controls respectively. (c) From 3288 alignments, 2608 showed a larger 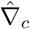 in *D. melanogaster* than *D. simulans*.

The magnitude of NMD was systematically elevated in *D. melanogaster* compared to *D. simulans*. The rightward shifted distribution of 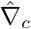 differences (Figure 3c) demonstrates that, on average, loci in *D. melanogaster* exhibit higher levels of NMD than their *D. simulans* ortholog.

### The Magnitude Of NMD Is Influenced By Purifying Selection In *Drosophila* Lineages

We observed that the magnitude of NMD was significantly elevated for X-linked loci compared to autosomes in the *Drosophila* lineages (Figure S6). This supports the hypothesized increase in NMD resulting from the higher level of purifying selection of the X chromosome. The median 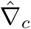 for Chromosomes 2, 3 and X in *D. melanogaster* were 0.0132, 0.0131 and 0.0177 respectively. The median 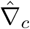 for Chromosomes 2, 3 and X in *D. simulans* were 0.0031, 0.0031 and 0.0040 respectively. Within a species, an unpaired *t*-test was performed between chromosomes (e.g., Chromosome 2 and X in *D. melanogaster*). All comparisons between an autosome and the X chromosome of the same species were significant, however, comparisons between the autosomes were not.

### Evidence for substantial NMD in the Human Genome

Our analysis of great ape genomes directly evaluated the impact of selection on NMD. We observed clear differences in the levels and magnitude of NMD between CDS and their adjacent intronic regions within the human genome (Figure 4). Using the TOE, the estimated proportion of data exhibiting NMD was 0.64 for CDS and 0.25 for intronic sequences, indicating the prevalence of NMD is more than two-fold greater in coding regions (Figure 4a-b). Further, comparisons of the magnitude of NMD confirmed that introns are closer to mutation equilibrium than their paired CDS regions. Across the evaluated loci, the histogram of paired differences in 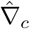 between CDS and introns 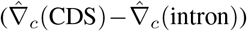 showed a rightward shift, reflecting systematically higher magnitude of NMD in coding regions (Figure 4c). The values for 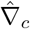 linked CDS and intron were significantly correlated according to the Spearman ranked-order test 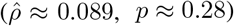.

**Figure 4.**
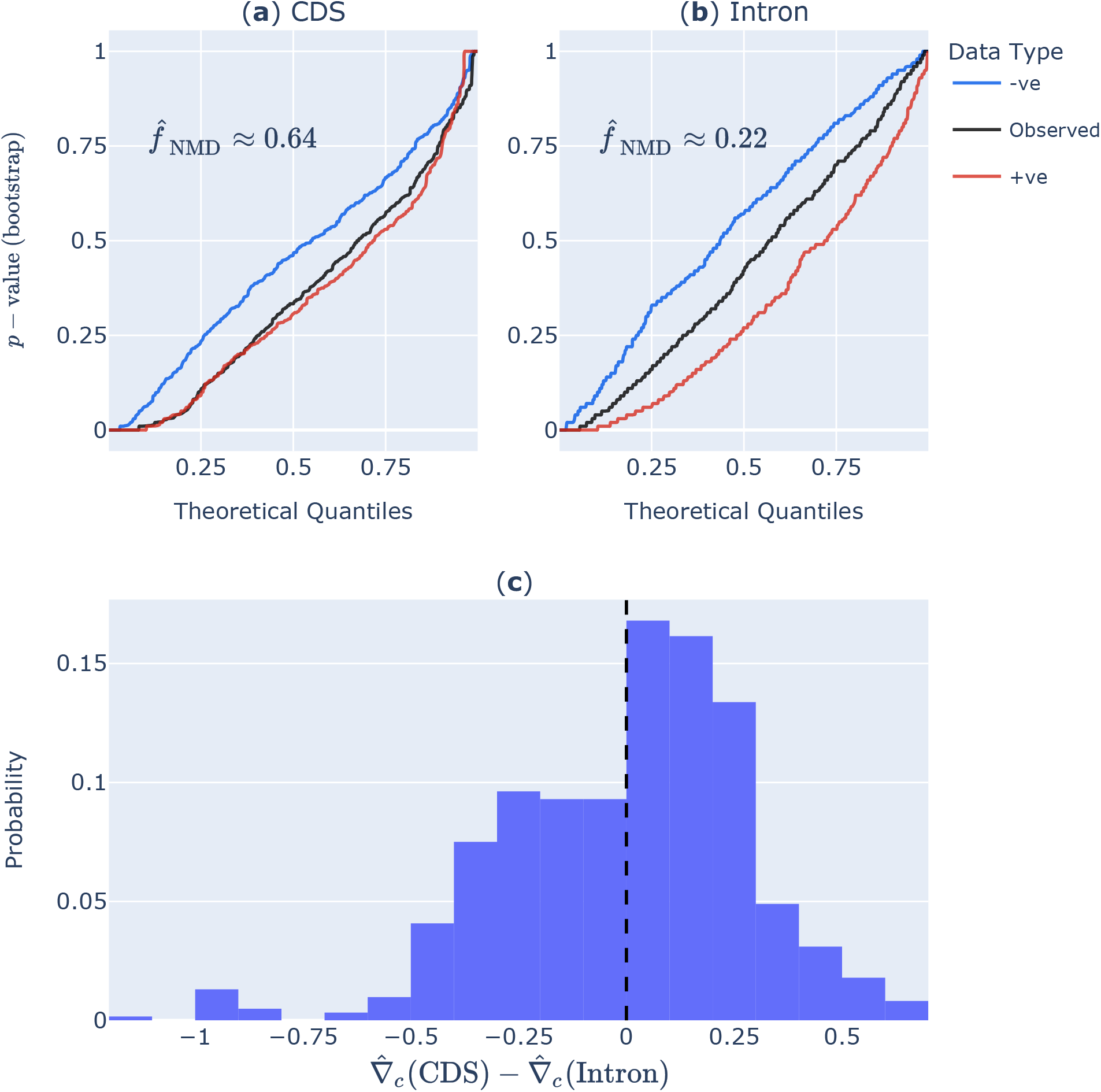
Strong support for greater NMD in Human CDS compared to flanking introns. Smile plots for (a) CDS and (b) Introns, indicated CDS were closer to the positive controls of non-stationary evolution than their intronic counterparts. (c) From 613 loci with paired CDS-Intron alignments, 349 showed a larger 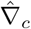 in the CDS than Intron.

## Discussion

We evaluated the widely employed assumption that sequence divergence processes are at equilibrium through the development and application of statistical procedures across distinct genomic and evolutionary contexts. Using synthetic data we established our statistical measures conform with theoretical expectations across a range of conditions that we expect cover most experimental contexts. The robust performance of the statistical measures was further demonstrated by application to empirical “knowns”. As expected, these datasets exhibited pronounced neutral mutation disequilibrium (NMD), confirming our prior predictions, and illustrating that our developed statistical framework is well equipped to detect and measure NMD. We also demonstrated that NMD is prevalent in the human genome. We further examine the implications of prevalent NMD, identifying the fields of interpretation of within and between species genetic variation where our current understanding requires revisiting.

To achieve consistency with theoretical expectations under the null model it was necessary to employ a parametric bootstrap. The structure of our Test Of the Existence of NMD (TOE) is such that the null nests within the alternate. Although asymptotic theory indicates the resulting Likelihood Ratio Test (LRT) statistic should be *χ*^2^ distributed in data that is consistent with the null, we observed deviations (Figure 2a) similar to what has been reported in other cases (e.g. Ota et al. 2000). A comparison with the test of Squartini and Arndt (2008) confirmed that its test statistic, originally assumed to be *χ*^2^ distributed, showed even more pronounced deviations (Figure S5). We suggest that for the TOE, this tendency for elevated false positives plausibly arises from overfitting as a result of the large number of free parameters. This conclusion is supported by the convergence towards the asymptotic expectation with increasing data (Figure 2a).

Results from our methods were consistent with prior predictions for the empirical datasets with known NMD. In the absence of an explicit expected result for natural data, we drew on predictions about inequalities between the region or lineage subjected to mutagenic changes and flanking genomic regions or sister lineages known to be unaffected. Furthermore, we explicitly generated synthetic negative and positive controls to establish the absolute boundaries attainable for the data sets, within which we examined the consistency of the observed results with our predictions. Variations in these boundaries across datasets highlighted that there exist inherent properties of natural data that must be considered in the interpretation of the statistics.

Although the limited precision of bootstrapped 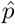-values prevented us from designating individual alignments as significantly in NMD, we were nevertheless able to estimate the overall fraction of such alignments. High-precision *p*-values are required for conventional multiple-hypothesis correction, and obtaining them through bootstraps would demand a prohibitive number of replicates. Instead, we adopted the procedure of Storey and Tibshirani (2003), which allowed us to estimate the proportion of alignments consistent with the alternate hypothesis.

Our analysis of *Fxy* demonstrated that our methods could accurately detect localized occurrences of NMD. The TOE detected significant NMD (at the *α* = 0.01 threshold) in all PAR-linked introns but not in the single X-linked intron, with all PAR-linked introns also exhibiting substantially higher levels of NMD (Table 1). Although this comparison depended on a single X-linked intron, it plausibly reflects real biological differences. As the X-linked intron is of comparable length to the longest PAR-linked intron, its non-significant TOE result is unlikely attributable to insufficient power. The differences in NMD magnitude are also unlikely driven by variations in alignment length, as our simulation study showed that the expected value of ∇_*c*_ is robust to such variability (Figure 2b). Although the simulation study also showed that the variance of ∇_*c*_ was inversely proportional to alignment length, the estimated standard deviations for the *Fxy* data were sufficiently small to support confidence in the measured values. The consistency in 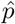-values from the TOE and 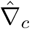 further provide confidence that the difference between these two groups of alignments truly reflects underlying NMD differences and thus the robust performance of our statistical measures.

Our analysis of *Drosophila* genomes demonstrated that our statistical methods could identify a genome wide elevation of NMD, as indicated by the systematic bias towards increased NMD in *D. melanogaster* (Figure 3a-c). Although the trend was not universal, as a small proportion of *D. melanogaster* orthologs had lower estimated NMD than their *D. simulans* counterparts (Figure 3c), these exceptions likely stem from sampling error or lineage-specific pressures on mutation. Nevertheless, the strong overarching pattern supports our prediction that the recent evolution of an anti-mutator allele in *D. melanogaster* has given rise to an elevated level of NMD.

We detected evidence that changes in natural selection may shape NMD signals, and thus plausibly contribute to the small discordance with predictions discussed above. As the equilibrium nucleotide frequency for a sequence is governed by the relationship between the mutagenic process and selection pressure, NMD can also arise due to changes in natural selection. In the *Drosophila* lineages, we interrogated this possibility by contrasting the NMD levels on X-linked vs autosomal chromosomes, given the X-linked genes are hemizygous in *Drosophila* males and are thus subjected to stronger purifying natural selection. Stronger purifying selection implies a slower rate of substitution and in turn, a slower rate of NMD decay. Thus we expected higher levels of NMD for X-linked loci. The X-linked genes were indeed systematically higher in their levels of NMD (Figure S6). These findings suggest that lineage-specific fluctuations in natural selection could contribute to the observed discrepancies in Figure 3c.

Our findings on the impact of selection from the *Drosophila* lineages also emerge in the Great Ape analyses. Consistent with the prediction that purifying selection acts more strongly on coding sequences (CDS) than on flanking introns, we observed both a higher fraction of CDS exhibiting NMD existence and a greater average magnitude of NMD signal in CDS compared to introns (Figure 4). However, although the fraction of NMD existence for CDS was approximately two times greater than that of introns, the difference in NMD magnitude was relatively weak (Figure 4c). This likely reflects the greater variance in estimates of 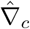 from the shorter CDS sequences, compounded with the limited number of substitutions that have accumulated between human and chimpanzee lineages. For this reason, we consider the fraction of NMD as a more reliable estimate due to the equivalence of sampling error between the sequence types.

For the Great Ape analyses, we had no prior hypothesis regarding the mechanism underpinning the NMD detected for the Human lineage. The marked difference in the proportion of encoded information between CDS (where a majority of nucleotides affect the encoded protein) and Intron sequence (where a minority of nucleotides encode regulatory sequence) leads to strikingly different exposure to natural selection. This makes it unlikely that natural selection would operate consistently across adjacent exon and intron sequences. In contrast, changes to processes affecting mutation (e.g. transcription coupled DNA repair, Svejstrup 2002) can feasibly influence co-transcribed exon and intron sequences. We did detect a significant positive correlation between 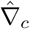 between CDS and introns. While the magnitude of the correlation was modest, this seems plausibly due to lower statistical power due to the factors discussed above. We therefore suggest the Human NMD is also due to changes in mutation.

### Implications

Our conclusions about the consistency of the statistics relied on comparisons between lineages or genomic regions with strong evidence for recent changes in mutagenesis and analogous samples known to be unaffected. It is noteworthy that even in those baseline samples, along with the protein coding component of the human genome, there were appreciable levels of NMD. In other words, NMD is not rare. So what are the implications of this?

The prevalence of NMD that we demonstrate calls into question the biological attribution assigned for many phenomena whose basis arises from application of time-reversible models. It has already been shown that using a time-reversible model to examine the molecular clock can lead to incorrect inference in terms of the existence and the nature of departure from the clock (Kaehler, Yap, Zhang, et al. 2015). It has likewise been demonstrated that failure to account for the non-stationarity of sequence evolution can lead to misinterpretation of the signal of natural selection when using codon models of sequence evolution (Kaehler, Yap, and Huttley 2017). The observation that stark differences in the existence and magnitude of NMD can exist between even adjacent nucleotides indicates that the common application of rate variation as an index of functional importance (as discussed by Graur et al. 2013) may be misattribution.

For analyses seeking to understand mechanisms underpinning sequence evolution it is clear that a non-stationary model, such as those available in cogent3 (Huttley, Caley, et al. 2025), should be applied. Even when the true divergence process is actually at NME, a more general model will give the same estimates (Kaehler, Yap, Zhang, et al. 2015). However, if the process is in NMD then only a more general model can sensibly be used. Failure to do so makes one vulnerable to misattributing mechanistic origins to patterns that may be merely artifacts of a model that is both wrong and misleading.

## Material And Methods

### Test of Existence of Neutral Mutation Disequilibrium

To formally assess whether the substitution process along a given branch exhibits Neutral Mutation Disequilibrium (NMD), we designed a Likelihood Ratio Test (LRT), termed the Test of Existence (TOE). As shown in Figure 1a, we designate the branch of interest the foreground branch, and all others branches the background branches (Figure 1). To test for disequilibrium in a single branch we model a continuous-time process on the foreground branch and discrete-time processes for the background branches, changing only the process on the foreground branch between the null and alternate models.

For the nominated background branches, the Barry and Hartigan (BH, 1987) discrete-time Markov process for nucleotides was used. We expect the root of the tree to occur on one of the background branches, at which point the direction of time will swap. Using BH avoids this potential confounder, as well as changes in mutagenesis on the background branches (Verbyla et al. 2013). Under the alternate model, the foreground branch follows a General Nucleotide process (GN, Kaehler, Yap, Zhang, et al. 2015). GN is a non-stationary and non-reversible nucleotide model with 12 distinct parameters in *Q* (note that GN and BH are identical when the BH process is embeddable within GN, Verbyla et al. 2013). Under the null model, the foreground branch follows the General Nucleotide Stationary process (GNS, Yap and Speed 2005). GNS is a Markov process that differs from GN by its constraint to being stationary with respect to a specified nucleotide frequency vector *π*. In our case, this is the nucleotide frequencies in the unobserved ancestor, which we denote *π*_0_. The constraint of stationarity is parameterized via the parameters of *Q*, in which GNS has 9 free parameters (Yap and Speed 2005).

The TOE is a likelihood ratio test (LRT) between the following two hypotheses: **Null**: the foreground evolved according to the **GNS**, the background according to BH; **Alt**: the foreground evolved according to the **GN**, the background according to BH. For this LRT, a rejection of the null means that the foreground branch was described better with a non-stationary process. Such a result suggests NMD precisely in the foreground branch. All TOE model fits assumed mixed discrete- and continuous-time Markov processes. For brevity, we will refer to a model by the process assumed on the foreground branch.

### Measuring the Magnitude of Neutral Mutation Disequilibrium

To quantify the magnitude of NMD, we introduced a novel statistic ∇, which measures the rate at which nucleotide frequencies have been converging toward equilibrium (Figure 1). Consider a process operating on a single branch of a phylogenetic tree for time interval *t*, with *π*_0_ the frequencies of nucleotides at the root, and the rate matrix *Q*. The nucleotide distribution at *t* is *π*(*t*) = *π*_0_ · *e*^*Qt*^. Under weak assumptions, a non-stationary process will converge to a stationary process, for which *π* remains unchanged over time. ∇ is defined as the Euclidean norm of the instantaneous rate of change of nucleotide frequencies.

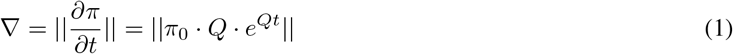

By virtue of its definition, 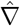 will be strictly non-negative and thus we expect it to exhibit a positive bias with sampling error. Given a truly stationary process and infinite data, we expect 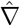 to approach 0. We note here that in our study design, we compare sister taxa or sequences on the same lineage. In these cases, we define time *t* = 1 thus allowing comparisons between 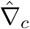.

### Parametric Bootstrapping

We applied parametric bootstrapping to address biases arising from limited sample sizes. As shown in Figure 2, both the TOE and ∇ were shown to be sensitive to the length of an alignment.

When asymptotic assumptions are shown not to hold, a robust estimate of the *p*-value corresponding to an LRT statistic can be derived from a null distribution generated parametrically (Goldman 1993). Using parameter estimates of the TOE null model fitted to an observed alignment, we generated the null distribution of LRT. (Due to the computational time required to fit both null and alternate models to an alignment we limited the number of synthetic alignments to 100.) The *p*-value was estimated by comparing the LRT statistic obtained from the observed alignment to this synthetic null distribution.

Alignment length also had a pronounced impact on ∇, with shorter alignments exhibiting a tendency for inflating ∇ on stationary processes. To correct for this bias, we define ∇_*c*_ as

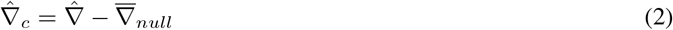

where 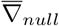 is the mean of ∇ estimated from synthetic alignments generated under the null. For practical reasons, we used the same alignments as generated for the hypothesis test parametric bootstrap.

### Data Sets

#### Microbial

We used a Microbial data set as the basis for our simulation study and further for evaluating the question concerning the prevalence of NMD in microbes. 16S ribosomal RNA (rRNA) sequences from the GreneGenes database (McDonald et al. 2012), as originally sampled by Kaehler, Yap, Zhang, et al. (2015), were downloaded from Dryad (https://doi.org/10.5061/dryad.g7g0n). The data consisted of 9,702 alignments of species triples that were sampled with an approximately uniform representation of maximum Jensen-Shannon Divergence (JSD). Increasing values of JSD indicate increasing differences in sequence composition between the sequences.

#### Drosophila

A *Drosophila* data set was applied to examine the influence of a mutator on NMD at a genomic scale. The selected data consisted of protein coding sequence (CDS) alignments of one-to-one orthologs from *Drosophila melanogaster, D. simulans* and *D. yakubra*. There were 9,237 alignments of one-to-one orthologs obtained from *flyDIVaS* (Stanley and Kulathinal 2016; Drosophila 12 Genomes Consortium et al. 2007).

#### Rodent

The localized occurrence of NMD was evaluated by examining the impact of translocation to the PAR on the *Fxy* gene in *Mus musculus*. To contrast the impact of translocation to the PAR in *M. musculus*, we sampled one-to-one orthologs of *Fxy* from *M. spretus* and *Rattus norvegicus*, in which *Fxy* is X-linked. Intronic *Fxy* sequences for the three species were sampled from Ensembl release 104 using EnsemblDb3 (Huttley 2021a). The first nine introns of *Fxy* were aligned using the Cogent3 progressive nucleotide aligner (Knight et al. 2007) using the HKY85 substitution model (Hasegawa et al. 1985) with default settings. The quality of the alignment path was visually examined using dotplots (Figure S2). Introns 4, 5 and 6 and the first 4,000bp of Intron 2 were deemed as being high quality alignments and were selected for analyses. Intron 3 was excluded as the X/PAR boundary lies within it while the remaining intron alignments were excluded due to their poor quality.

#### Great Apes

Sampling CDS and introns from the same gene enabled paired comparison of the two sequence types. Data for the Great Apes intronic and CDS data sets were sampled from Ensembl release 104 (Howe et al. 2020). From *Homo sapiens* (human) protein coding genes of chromosome 1, we identified one-to-one orthologs in *Pan troglodytes* (chimpanzee) and *Gorilla gorilla* (gorilla) using EnsemblDb3 (Huttley 2021a) and homologsampler (Huttley 2021b). From this gene list we sampled the unaligned CDS from the canonical transcript (Howe et al. 2020). Intron alignments were obtained directly from whole genome alignments via the human gene coordinates, with exons filtered out resulting in an alignment of concatenated introns. We aligned CDS using the Cogent3 progressive codon aligner (Knight et al. 2007) using the MG94HKY (Muse and Gaut 1994) substitution model with default settings.

### Data Filtering

Substitution models were explicitly restricted to interchanges between nucleotide states, so alignment positions linked to other mutation types were removed. This included gap characters and annotated simple tandem repeats (likely to have evolved through strand slippage, Levinson and Gutman 1987). For all CDS, alignments shorter than 900bp were excluded. To reduce the impact of selection in CDS alignments, we then selected the third codon positions only. For intron alignments from the great apes, the minimum length was 3, 000bp. (See Table S1 for summary statistics).

### Experimental Design

#### Establishing properties of the statistical measures in synthetic data

We sought to establish the robustness of the developed methods by analysis of synthetic data which captures the extremes of two dimensions of sequence divergence pertinent to our study. Nucleotide frequencies are integral to the definition of NMD and we furthermore want to avoid confounding historic NMD, whereby sequences differ in their composition even though they have reached equilibrium. We therefore considered proxy measures of both of these properties and devised experiments to establish that our statistical procedures are robust to combinations of their extremes. The presence or absence of historic NMD between taxa was quantified using Jensen-Shannon divergence (JSD) between the in-group sequences. Balanced or imbalanced nucleotide frequency distribution was quantified using the Shannon Entropy of the GNS stationary distribution vector (*H*(*π*_∞_)). Four “seed” alignments (Figure S3) were selected from the Microbial data that represented the different combinations of Low or High JSD and Low or High Shannon’s entropy. (See Supplementary Materials for the full procedure.)

As the robustness of maximum-likelihood techniques is sensitive to alignment length (Ota et al. 2000) we generated synthetic data to facilitate assessing the influence of alignment length on our statistics. Alignment lengths chosen as representative of the biological applications were: 300bp, 3,000bp, and 30,000bp. Each simulation set consisted of 1,000 synthetic alignments generated using the null model of the TOE, derived from the parameter estimates from one seed alignment.

#### Empirical knowns

We have chosen the Drosophila and Rodent data sets as positive empirical controls, natural occurrences of the alternate hypotheses. In these cases we have a strong expectation of the existence of NMD based on putative perturbations to mutagenesis, leading to specific hypotheses about the expected form of NMD.

#### Simulated positive and negative controls

For the empirical data sets involving more than 100 alignments, we generated paired distributions of TOE *p*-values from the fitted null and alternate models. As an example, consider the Drosophila 5,680 CDS alignments. For a single *observed* alignment, the parameter estimates from the fitted null model were used to generate a single synthetic null alignment which was added as a negative control for that *observed* alignment. A synthetic alternate alignment generated by the alternate model was likewise added as a positive control for that *observed* alignment. Both collections of control alignments were analyzed using the TOE.

### Algorithmic Implementation and Data Availability

All methods were written in Python. For all analysis of biological sequence data, we used the develop branch of Cogent3 (Huttley, Caley, et al. 2025). The code implementing the statistical methods provides a command line tool (mdeq) for computing NMD. Explicit tests of code correctness were implemented covering 94% of all code lines. This tool can be installed from the Python Package index (https://pypi.org), with source code available at GitHub https://github.com/HuttleyLab/MutationDiseq and on Zenodo (DOI 10.5281/zenodo.16829613). All code for data sampling and application of mdeq is available in a GitHub repository https://github.com/HuttleyLab/ MutationDiseqAnalysis and on Zenodo (DOI 10.5281/zenodo.16830090). All data used and results produced in this study are available at DOI 10.5281/zenodo.16916309.

Analyses that required model fitting were performed on the National Computational Infrastructure’s (NCI, https://nci.org.au) supercomputer Gadi. Gadi uses a Linux operating system, and analyses were run on multiple nodes comprising 24-core Intel Xeon Scalable ‘Cascade Lake’ processors. All other analyses were performed on a MacBook Pro with four Intel i7 processors on which the operating system was MacOS version 11.4.

## Acknowledgments

This work was supported by the NCI Adapter Scheme, with computational resources provided by NCI Australia, an NCRIS-enabled capability supported by the Australian Government. We thank Eric Stone for comments on a draft of this work.

