## Supplementary figures and tables for "Neutral mutation disequilibrium is common in prokaryotes and animals"

### Supplementary material for: *Neutral mutation disequilibrium is common in prokaryotes and animals*

#### Supplementary Background

##### Evolutionary History of Fxy in Rodents

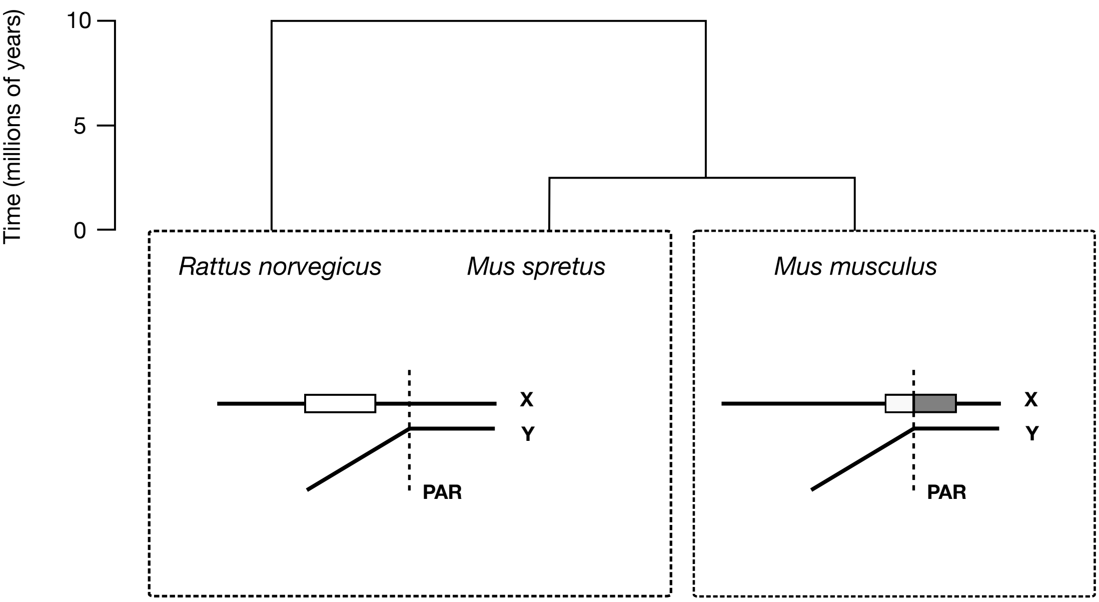


Figure S1: **Evolutionary History of *Fxy* in Rodents**. The *Fxy* gene in *M. musculus* was translocated from a X-specific position to a new position in which it overlaps with the Pseudo-Autosomal Region (PAR). The overlap in *M. musculus* is shown as the shaded region of the gene, with the boundary to the PAR falling in intron 3 [Palmer et al., 1997]. The divergence timescale is indicated in millions of years. In rodents and other mammals, *Fxy* is X-specific, suggesting translocation to the PAR in *M. musculus* after divergence with *M. spretus*, between 2-3 million years ago [figure adapted from Figure 1 of Galtier and Duret, 2007].

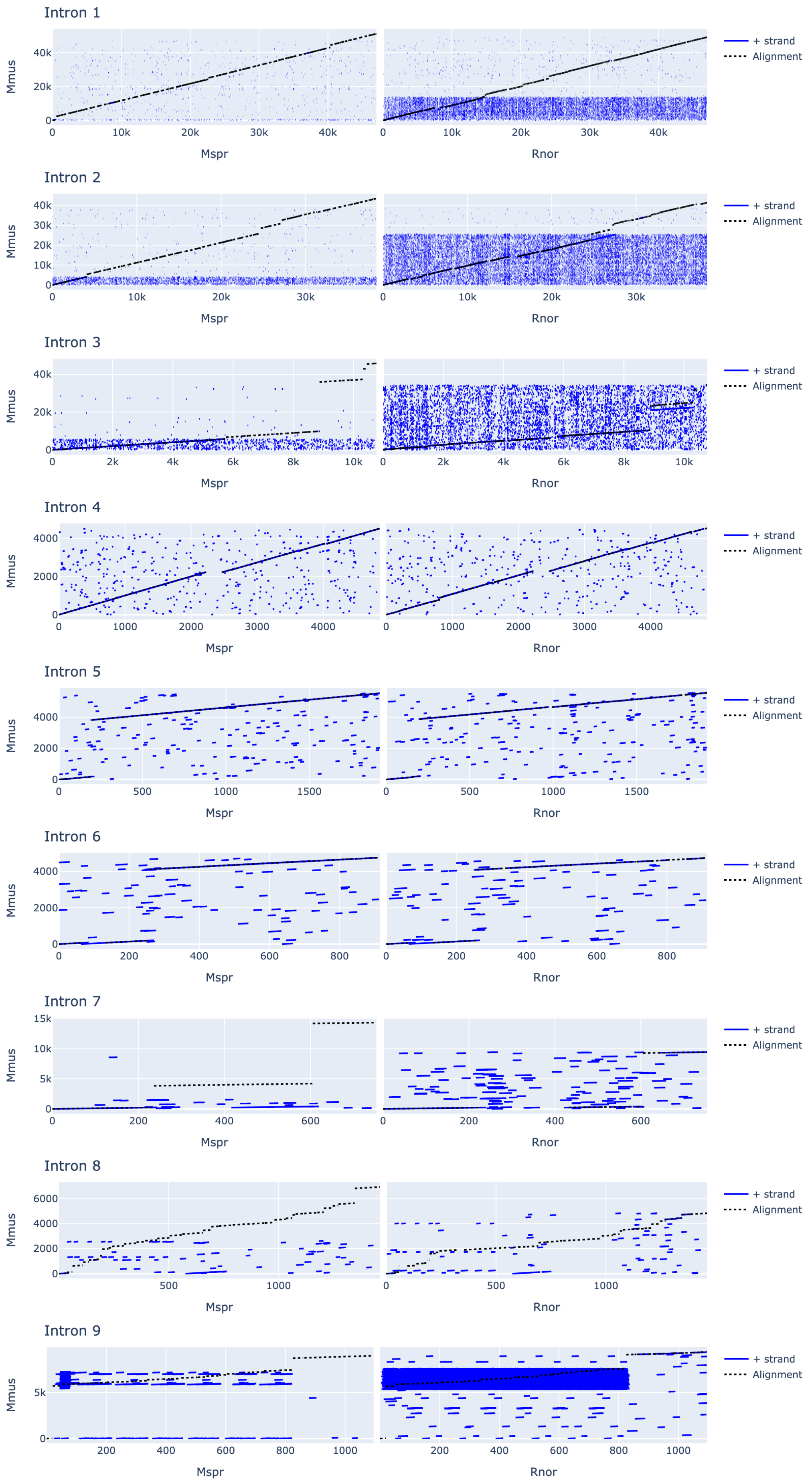


Figure S2: Dotplots of $F_{xy}$ introns. Mmus - *M. musculus*, Mspr - *M. spretus* and Rnor - *R. norvegicus*. The solid line represents runs of matching sequence while the dashed line is the path found by the alignment algorithm. An alignment path that does not overlay a diagonal blue line indicates a poor alignment.

### Supplementary Methods

#### Data sampling

##### Summary of the data sets

| Data Set | Bio Type | min(length) | median(length) | max(length) | # |
|---|---|---|---|---|---|
| Microbial | 16S rRNA | 954 | 1,140 | 1,275 | 380 |
| Great Apes | Intron | 3,038 | 13,561 | 251,535 | 613 |
| Great Apes | CDS | 300 | 576 | 3,773 | 613 |
| Rodent | Intron | 834 | 2,701 | 4,358 | 4 |
| Drosophila | CDS | 300 | 533 | 8,892 | 5,680 |

Table S1: Summary statistics of filtered data sets. Range of alignment lengths and the total number of alignments are shown. # is the number of alignments

##### TOE – Test of Existence

The Test of Existence of equilibrium (TOE) examines the nature of the substitution process operating on a designated foreground branch (see main manuscript Figure). Under the null model, substitution on the foreground branch is modeled using a non-reversible but stationary substitution model (referred to as the General Stationary Nucleotide model, GSN). Under the alternate, the foreground branch is modeled using a non-reversible and non-stationary substitution model (referred to as General Nucleotide, GN, Kaehler et al. [2015]). In both the null and alternate, the background branches are modeled using a discrete-time Markov process, which corresponds to the Barry-Hartigan (BH) substitution model Barry and Hartigan [1987]. The BH is the most general possible nucleotide substitution model Verbyla et al. [2013].

##### Maximum Likelihood

We employed the phylogeny-based maximum likelihood (ML) framework [Felsenstein, 1981] for statistical inference. A likelihood ratio test (LRT) statistic is the ratio of the log-likelihoods of the two models:

$$\mathrm{LR} = 2(\ln \mathcal{L}_{alt} - \ln \mathcal{L}_{null}).$$

If the null model nests within the alternate the LRT statistic can be distributed $\chi^2_{\mathrm{df}}$ with degrees freedom (df) equal to the difference in the number of free parameters between the models

[Kendall and Stuart, 1979]. For a given alignment, whether this property holds is dependent on the alignment length. When this condition is not satisfied, a parametric bootstrap [Goldman, 1993] was employed to estimate the $p$-value.

Under mild conditions ML is consistent, meaning that as the amount of data (the length of the alignment) tends to infinity, the probability of obtaining the true value of the parameters tends to one [Chang, 1996]. Salient conditions required are that the alignment contains at least three sequences [Chang, 1996], that the P matrices satisfy the diagonal largest-in-column property [Chang, 1996] and that there is a unique mapping of Q $\rightarrow$ P [Kaehler et al., 2015]. For these reasons all data sampling ensured data sets contained three sequences and fitted models were checked to ensure they satisfied the remaining conditions.

#### Evaluation of model properties using simulated data

##### Selection of seed alignments for the simulation study

We fit a time-heterogeneous GN model to each alignment in the Microbial data set. For each alignment, we automatically selected a branch as the "foreground" (see the main manuscript) via the following procedure. We first identify the sequence pair with the smallest pairwise genetic distance as the in-group sequences. The in-group sequence with the largest average Jensen-Shannon divergence (JSD) was selected as the foreground.

We excluded from further consideration any alignment if:

- any maximum-likelihood estimates (MLEs) were within machine precision of the lower (1e-5) or upper (1e2) bounds
- the foreground matrix had a condition number $> 2$ [Schranz et al., 2008]

After applying this procedure, 1,779 alignments remained. From these, we estimated the magnitude of non-stationarity using $\widehat{\text{JSD}}$ between the inferred in-group sequences (those with the pairwise shortest genetic distance). We further estimated the stationary distribution of the foreground matrix and computed the Shannon Entropy of this vector ($\hat{H}(\pi_\infty)$). The latter statistic is indicative of the evenness of the nucleotide frequency distribution.

The selected 4 seed model fits (Fig S3) were for alignments: 197113_332182_17210, 19 8257_206396_13724, 200580_114946_573911, 758_443154_73021 and their corresponding designations are shown in Table S2.

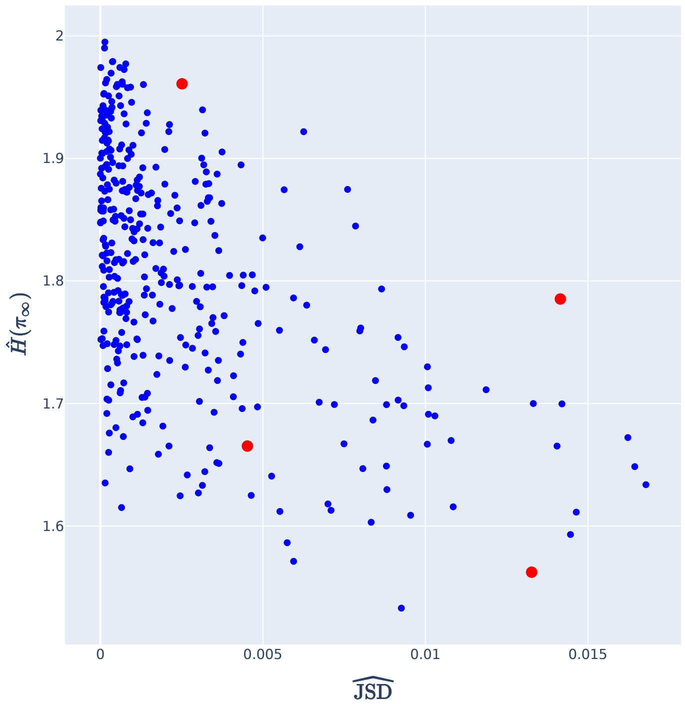


Figure S3: Sampled seed alignments. The distribution of $\widehat{JSD}$ by Shannon's entropy of the equilibrium nucleotide distribution ($\hat{H}(\pi_\infty)$ was used to identify 4 alignments from the branches. The red markers correspond to the Entropy × JSD model fits selected for the simulation study and correspond to the (clock-wise from top left) Hi-Lo, Hi-Hi, Lo-Hi, Lo-Lo (Entropy-JSD) conditions.

| Identifier | Entropy | JSD |
|---|---|---|
| 197113_332182_17210 | Hi | Hi |
| 198257_206396_13724 | Hi | Lo |
| 200580_114946_573911 | Lo | Hi |
| 758_443154_73021 | Lo | Lo |

Table S2: Selected seed fits from microbial data. Fits from these alignments were used for the simulation study. Identifier is from the GreenGenes alignment. Entropy and JSD categories are from Figure S3.

**Simulation of data for evaluating statistical performance of the Test of Existence (TOE)**

For each of the seed alignments, we used the MLEs from the observed alignment with the exception of the state frequencies of the unobserved ancestor ($\pi_0$). We ensure evolution on the foreground branch is consistent with the null model of equilibrium by setting $\pi_0 = \pi_\infty$(foreground). Synthetic alignments were generated under this parameter set using standard `cogent3` capabilities.

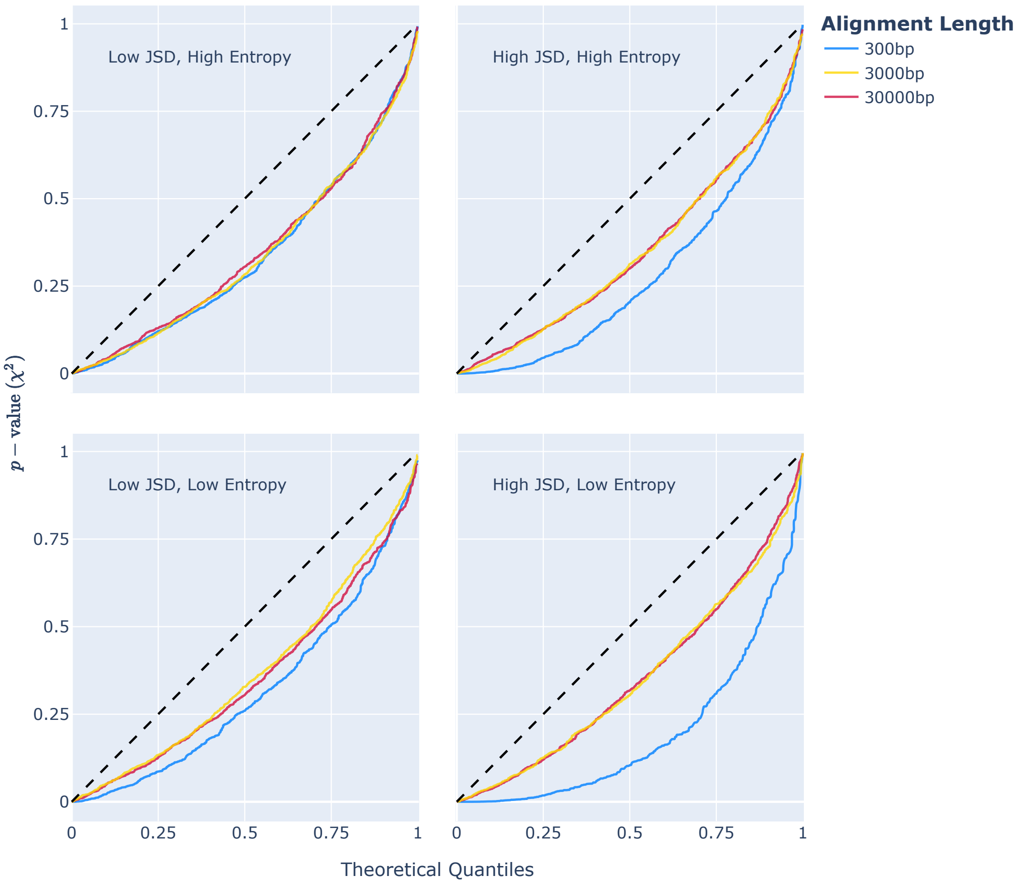


Figure S4: TOE $p$-values from the theoretical distribution are not uniform. Subplots are from seed alignments simulated under the null corresponding to the Entropy-JSD conditions.

### Supplementary Results

#### TOE likelihood ratio test statistics are inconsistent with the $\chi^2$ distribution

The quantile plots indicate that the test is strongly biased towards overestimation of significance for all alignment lengths and all conditions of entropy and JSD examined (Figure S4).

#### The test statistic presented in Squartini and Arndt [2008] was not consistent with the assumed theoretical distribution

The quantile plots indicate that the $\chi^2$ test presented by Squartini and Arndt [2008] was extremely biased towards overestimation of significance for all alignment lengths and all conditions of entropy and JSD examined (Figure S5). We note here that our implementation of this test was a variant of that described by Squartini and Arndt. Their original expression defined the *observed* frequency vector from an entire alignment. We needed to modify this test since our

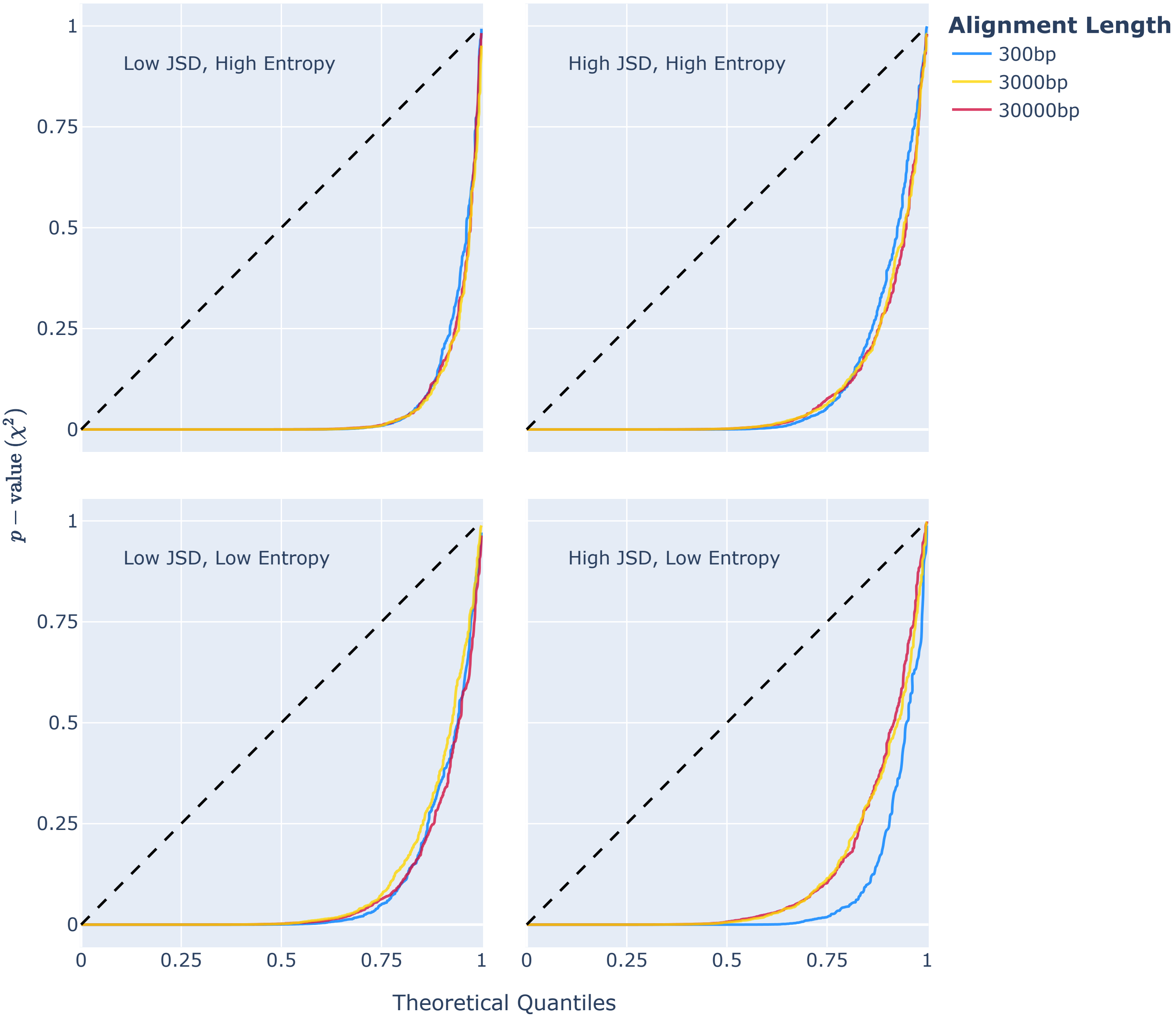


Figure S5: A test statistic derived from Squartini and Arndt [2008] was not consistent with the assumed theoretical distribution. The expected relationship is shown as the dashed line. Subplots are from seed alignments simulated under the null corresponding to the Entropy-JSD conditions.

simulation allowed the background edges to be undergoing non-stationary evolution. We thus defined the frequency vector from the foreground sequence as our *observed*. The results from this are displayed in Figure S5. Note that departure from theoretical expectation was worse under the original Squartini and Arndt [2008] definition (result not shown).

#### Genomic distribution of $\hat{\nabla}_c$ in Drosophila

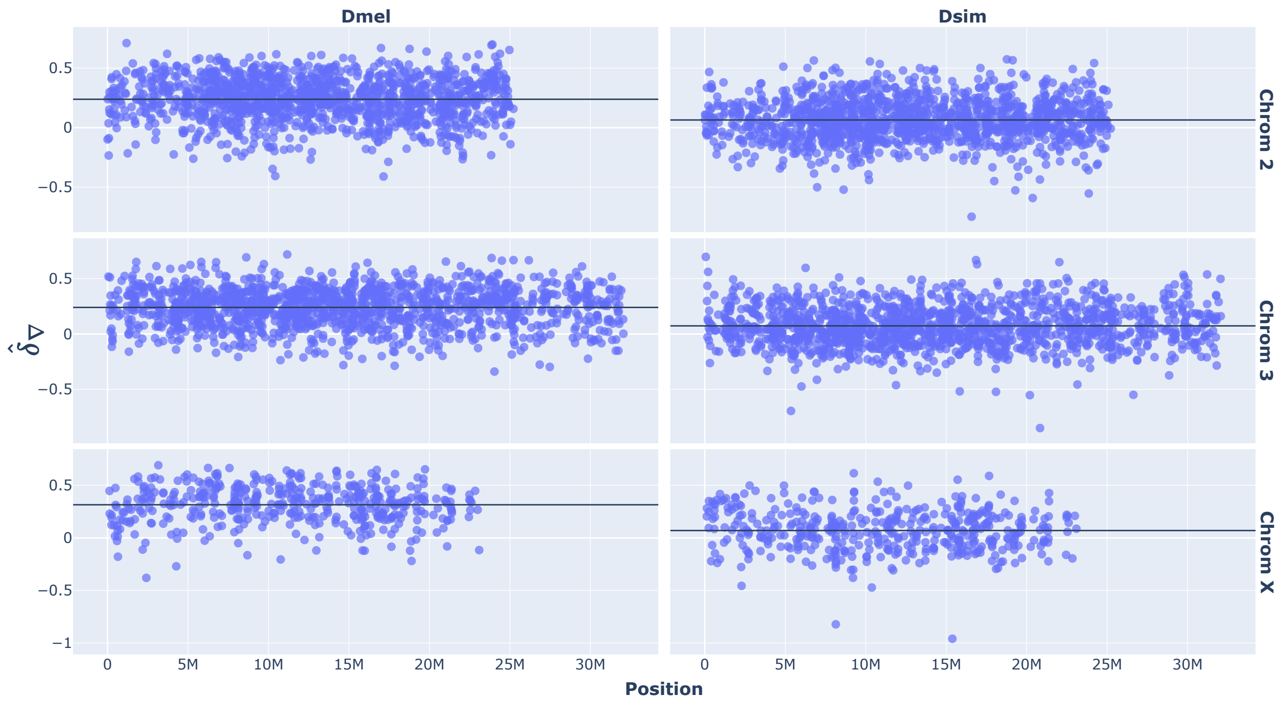


Figure S6: The magnitude of mutation disequilibrium is systematically larger across the genome of *D. melanogaster* (Dmel) compared to *D. simulans* (Dsim). The genomic coordinate of the gene in *D. melanogaster* was used for both Dmel and Dsim. The horizontal black line is the mean of the distribution for the chromosome indicated on the right-hand side of the figure.
